# Acquired interbacterial defense systems protect against interspecies antagonism in the human gut microbiome

**DOI:** 10.1101/471110

**Authors:** Benjamin D. Ross, Adrian J. Verster, Matthew C. Radey, Danica T. Schmidtke, Christopher E. Pope, Lucas R. Hoffman, Adeline M. Hajjar, S. Brook Peterson, Elhanan Borenstein, Joseph D. Mougous

**Author notes:** Equal contribution (Benjamin D. Ross, Adrian J. Verster).

## Abstract

The impact of direct interactions between co-resident microbes on microbiome composition is not well understood. Here we report the occurrence of acquired interbacterial defense (AID) gene clusters in bacterial residents of the human gut microbiome. These clusters encode arrays of immunity genes that protect against type VI secretion toxin-mediated intra- and inter-species bacterial antagonism. Moreover, the clusters reside on mobile elements and we demonstrate that their transfer is sufficient to confer toxin resistance *in vitro* and in gnotobiotic mice. Finally, we identify and validate the protective capacity of a recombinase-associated AID subtype (*r*AID-1) present broadly in Bacteroidales genomes. These *r*AID-1 gene clusters have a structure suggestive of active gene acquisition and include predicted immunity factors of toxins deriving from diverse organisms. Our data suggest that neutralization of contact-dependent interbacterial antagonism via AID systems shapes human gut microbiome ecology.

---

Polymicrobial environments contain a plethora of biotic and abiotic threats to their inhabitants. Bacterial survival in these settings necessitates elaborate defensive mechanisms. Some of these are basal and protect against a wide range of threats, whereas others, for instance CRISPR-Cas, represent adaptations unique to the specific threats encountered by a bacterial lineage (*1–3*). The density of bacteria in the mammalian gut microbiome can exceed 10^11^ gm^−1^; therefore, overcoming contact-dependent interbacterial antagonism is likely a major hurdle to survival in this ecosystem (*4, 5*). The type VI secretion system (T6SS) is a pathway predicted to be widely utilized by gut bacteria to mediate the delivery of toxic effector proteins to neighboring cells (*6–9*). While kin cells are innately resistant to these effectors via cognate immunity proteins, it is unknown whether non-self cells in the gut can escape intoxication.

To identify potential mechanisms of defense against T6S-delivered interbacterial effectors, we mined a large collection of shotgun metagenomic samples (n=553) from multiple studies of the human gut microbiome for sequences homologous to known immunity genes (table S1) (*10–12*). We first focused our efforts on *Bacteroides fragilis,* a bacterium belonging to the order Bacteroidales, which harbors a well described and diverse repertoire of effector and cognate immunity genes (table S2) (*6, 7, 13*). As expected for genes predicted to reside within the *B. fragilis* genome, sequences mapping to these immunity loci were detected at an abundance similar to that of *B. fragilis* species-specific marker genes in many microbiome samples (Fig. 1A, grey; table S1). However, in a second subset of samples, immunity genes were detected at an abundance significantly higher (>10X) than expected given the abundance of *B. fragilis* (Fig. 1A, blue). Finally, we identified a third subset of samples wherein immunity gene sequences were detected in the absence of *B. fragilis* (Fig. 1A, green). These latter sequences include close homologs of 12 of 14 unique immunity genes (i1-i14) encoded by *B. fragilis* (Fig. 1B). In contrast to the pattern observed for immunity genes, we found no samples in which the abundance of corresponding cognate effector genes exceeded that of *B. fragilis* by a significant margin (Fig. 1C, table S1).

**Fig. 1.**
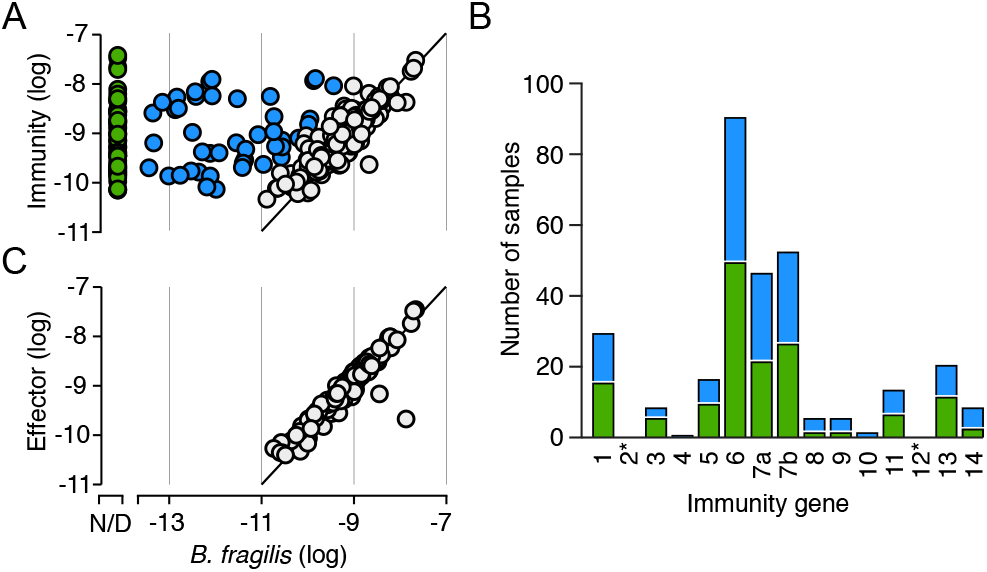
T6SS orphan immunity genes are found in in human gut microbiomes. (**A,C**) Comparison of the abundance of *B. fragilis*-specific T6SS immunity genes (**A**) or effector genes (**C**) with *B. fragilis* marker genes in adult microbiome samples derived from the HMP and MetaHIT studies (table S1). Abundance values denote the number of reads mapped to the gene, normalized by gene length and total number of reads in the sample. Immunity (or effector) gene abundances are calculated as the average abundance of all immunity (or effector) genes present in a sample. *B. fragilis* abundance is similarly calculated as the average abundance of all *B. fragilis* species-specific marker genes. Samples with undetectable *B. fragilis* (green) and samples in which immunity gene abundance exceeds that of *Bacteroides* by over 10-fold (blue) are highlighted. (**B**) The number of adult human gut microbiome samples in which the indicated immunity genes (1-14, GA3_i1-14 from ref (*6*)) can be detected at an 80% nucleotide identity threshold and at abundance over 10-fold higher than that of *B. fragilis* marker genes. Bars colored corresponding to the source of the immunity gene from (**A**) and asterisks indicate immunity genes without orphan representation.

The detection of *B. fragilis* immunity gene homologs in samples in which we were unable to detect *B. fragilis* strongly suggests that these elements are encoded by other bacteria in the gut. Indeed, BLAST searches revealed the genomes of several *Bacteroides* spp. that harbor *B. fragilis* T6S immunity gene homologs, including *B. ovatus* (i6, i7, i5, i14), *B. vulgatus* (i8, i13), *B. helcogenes* (i1, i9, i10), and *B. coprocola* (i8, i13). To determine whether these bacteria could also account for the presence of immunity genes in the human gut microbiome, we assembled full-length predicted immunity genes from the metagenomic sequencing reads of individual microbiomes. We limited this assembly to homologs of i6, the most prevalent immunity gene detected in samples lacking *B. fragilis* (Fig. 1B). Clustering of the recovered homologs showed that the majority of i6 sequences distribute into three discrete clades that differ by multiple nucleotide substitutions (i6:cI-cIII) (Fig. 2A and table S3). A comparison of these immunity sequences to available bacterial genomes revealed a clade matching cognate immunity genes in *B. fragilis* (i6:cI). Additionally, we found an i6 sequence homologous to i6:cIII in the genome of *B. ovatus,* which we previously found does not contain the cognate T6SS effector gene (*6*).

**Fig. 2.**
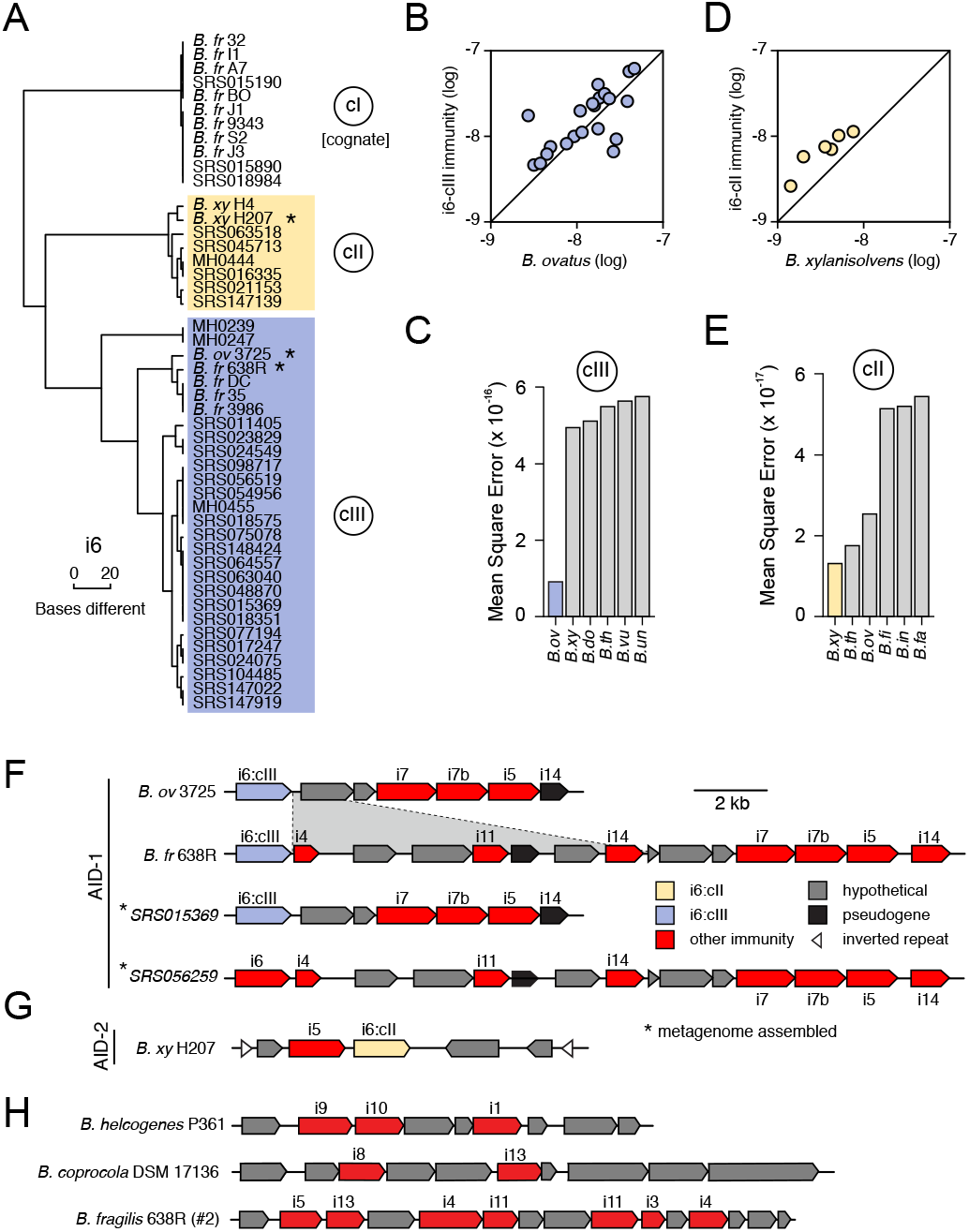
T6SS orphan immunity gene clusters are encoded by multiple species in the human microbiome. (**A**) Dendrogram depicting hierarchical clustering of orphan immunity gene i6 sequences extracted from genomes (*n* = 15) and metagenomes (*n* = 32). For genomes, immunity genes were identified using BLAST, while for metagenomes gene sequences were constructed from a pileup of shotgun reads aligned to the cognate reference sequence. Only samples with at least 10X coverage over 90% of the length of the gene were included. Sequence clades discussed in the text are denoted (cI-III) and those corresponding to orphan sequence are delineated with shaded boxes. Asterisks indicate strains for which orphan immunity gene clusters are shown in panels **F** and **G**. Strain name abbreviations defined in table S2. (**B-D**) Comparison of the abundance of genes from the indicated orphan immunity clades (colors as in (**A**)) with *B. ovatus* (B) or *B. xylanisolvens* (D) marker genes in adult microbiome samples derived from the HMP (SRS) and MetaHIT (MH) studies (table S1). (C,E) Linear model error values for the six species that best fit the i6:cIII (C) and i6:cII (E). Gene abundance calculated as in Fig. 1. (**F-H**) Depiction of representative AID-1 (**F**), AID-2 (**G**) and other (**H**) gene clusters containing homologs of the indicated *B. fragilis* T6S immunity genes. Sequences derived from genomes or metagenomic assemblies as indicated. The i6 gene of SRS056259 did not meet our sequence depth coverage requirements for inclusion in i6:cIII. The *B. xylanisolvens* strains in (**G**) was isolated and sequenced as a part of this study (BioProject PRJNA484981). Pseudogenes defined as genes with contiguous homology flanking nonsense codons. A region of difference between otherwise homologous clusters in *B. ovatus* 3725 and *B. fragilis* 638R is highlighted.

To identify the species encoding these sequences in human gut microbiomes, we used a simplified linear model to identify *Bacteroides* spp. whose abundance in microbiome samples best fits that of each immunity sequence clade. We found that i6:cIII is best explained in gut metagenomes by *B. ovatus* (Fig. 2B,C), suggesting that although reference genomes of both *B. fragilis* and *B. ovatus* contain these sequences, it is most often harbored by the latter in natural populations. We could not confidently define a single species harboring i6:cII by this method (Fig. 2D,E). Rather, multiple species exhibited similar error, a finding that is likely explained by the documented covariance of certain *Bacteroides* spp. in human microbiomes (*12*). To more conclusively define the origin of i6:cII genes, we leveraged metagenomic data and matching stool samples available from a recent study of the infant microbiome (*14*). Employing the same analysis pipeline utilized for the adult microbiomes, we identified several samples within this cohort containing i6:cII genes at an abundance in excess of *B. fragilis* (fig. S1 and table S4). Next, we isolated *Bacteroides* spp. from corresponding stool samples, identified candidate i6-positive strains by PCR and conducted whole genome sequencing, which defined the strains as *B. xylanisolvens*. Notably, the abundance of this same species best fit the abundance of i6:cII genes, albeit slightly, in the large adult metagenomic datasets (Fig. 2D). Based on these observations, we conclude that genes closely related to T6S cognate immunity genes of *B. fragilis* effectors are encoded by *B. fragilis* as well as other species of *Bacteroides* in human gut microbiomes. We hypothesize that these orphan immunity genes serve an adaptive role in the gut by providing defense during intra- and inter-species antagonism.

To gain insight into the function of orphan immunity genes, we examined their genomic context in available reference genomes. Using identity criteria similar to those employed in our metagenomic analyses, we found that homologs to *B. fragilis* T6S immunity genes i6 and i7 are located together within discrete gene clusters in several *Bacteroides* strains (Fig. 2F). The organization and nucleotide sequence of these clusters, which we termed AID-1 (acquired interbacterial defense 1) systems, is conserved between *Bacteroides* spp. Assembly of sequence scaffolds from metagenomic data confirmed the presence of AID-1 gene clusters in gut microbiome samples. Upon examination of AID-1, we identified distant homologs and pseudogenized remnants of several additional *B. fragilis* T6S immunity genes, including i4, i5, i11, and i14 (Fig. 2F and table S5). These findings prompted us to search reference genomes for more distant homologs of *B. fragilis* T6S immunity genes. In these searches, we additionally found gene clusters containing orphan homologs of i1, i3, i8, i9, i10, and i13 in *Bacteroides* genomes (Fig. 2H). Our identification of distant homologs to *B. fragilis* immunity genes within genomes was corroborated by querying a catalog of non-redundant complete gene sequences compiled from human microbiome samples, revealing related genes are found within gut bacteria (fig. S2). In *B. xylanisolvens*, we found that genes belonging to i6:cII are located in a unique, but analogous context adjacent to a homolog of i5 on an apparent transposable element (Fig. 2G) (*15*). We designate this sequence as the AID-2 system.

We next sought to define the phenotypic implications of orphan immunity genes of *Bacteroides* spp. during competition with *B. fragilis. B. fragilis* 9343 encodes the cognate effectors for i6 and i7, and prior data demonstrate that the corresponding toxins, e6 and e7, respectively, efficiently antagonize assorted *Bacteroides* spp. *in vitro* and in gnotobiotic mice (*7*). We thus employed this strain in growth competition assays against *Bacteroides* spp. bearing orphan immunity genes, derivative strains containing deletions of these genes, or genetically complemented strains. These experiments showed that in both *B. ovatus* and *B. fragilis*, AID-1 system genes grant immunity against corresponding T6S effectors (Fig. 3A, fig. S3A,B). The i6 and i7 genes of *B. ovatus* did not influence the outcome of its competition with *B. fragilis* 638R, which possesses an orthogonal effector repertoire (fig. S3C). Finally, we also found that an i6:cII gene from a *B. xylanisolvens* AID-2 system affords this bacterium protection against e6 of *B. fragilis* 9343 (Fig. 3B). In total, these data show that the orphan immunity genes of multiple *Bacteroides* spp. – localized to AID systems – can confer protection against effectors delivered by the T6SS of *B. fragilis*.

**Fig. 3.**
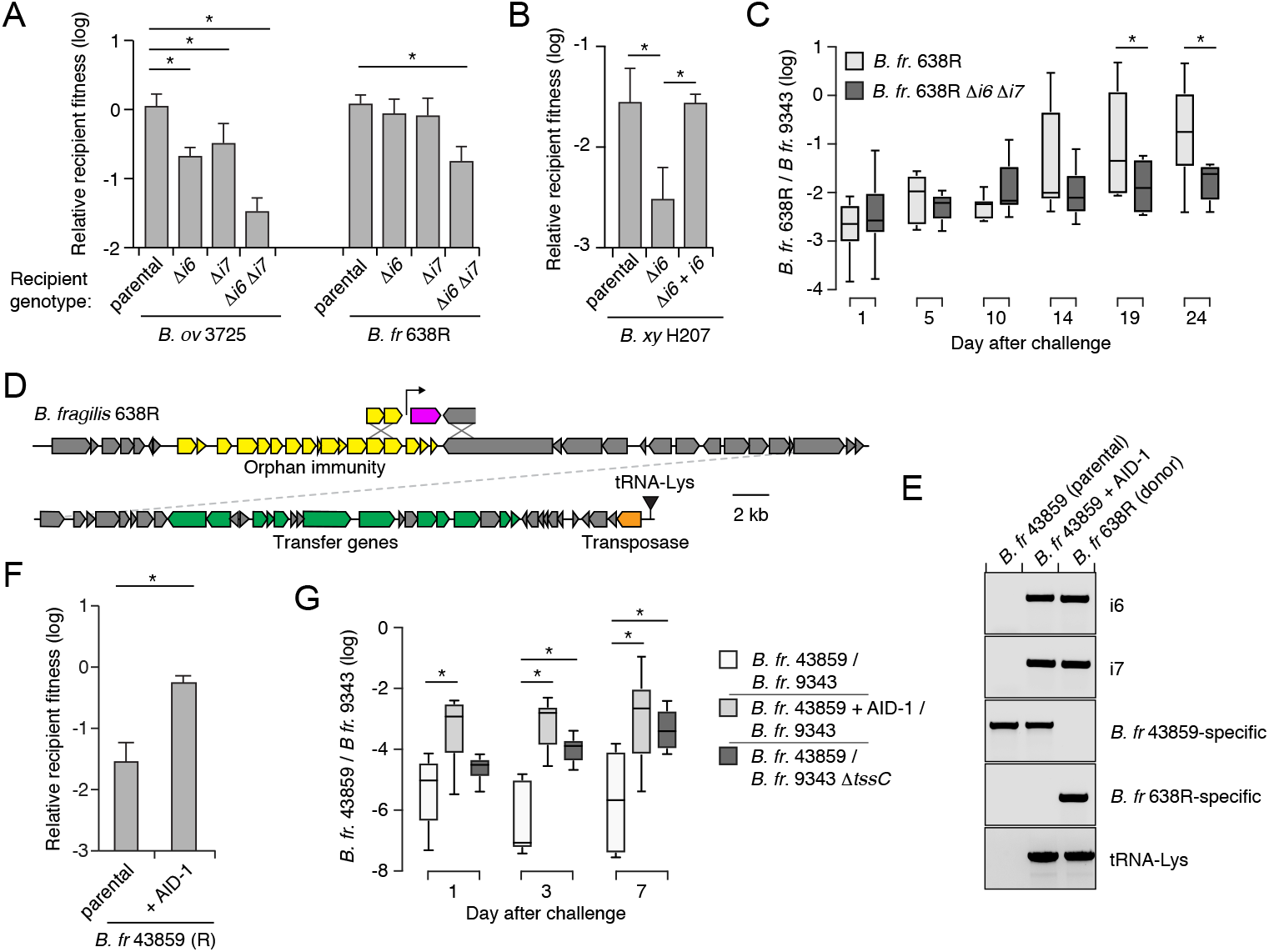
Orphan immunity genes are mobile and protect against T6S-delivered toxins *in vitro* and *in vivo.* (**A,B**) Outcomes of *in vitro* two-strain growth competition assays between the indicated AID-1 (*B. ovatus* 3725 and *B. fragilis* 638R (A)) or AID-2 (*B. xylanisolvens* H207 (**B**)) harboring strains versus *B. fragilis* NCTC 9343. Relative recipient fitness was determined by calculating the ratio of final to initial colony forming units (c.f.u.) and normalizing to the corresponding experiment with *B. fragilis* NCTC 9343 lacking *tssC* (T6S-inactive). Data represent mean ± s.d. of three replicates. Asterisks, *P* < 0.01 (unpaired Student’s t-test). (**C**) Outcome of pairwise competition between the indicated *B. fragilis* strains in germ-free mice. Mice were pre-colonized with *B. fragilis* 9343 for one week prior to challenge with 638R (N=8 mice per group in two independent experiments). Bacteria were enumerated by selective plating of fecal samples on the days indicated. Boxplots represent the interquartile range with indicated means for each group. Asterisk, *P* < 0.01 (Mann-Whitney U test for each time point). (**D**) Schematic depicting an ICE element from *B. fragilis* 638R harboring the AID-1 cluster depicted in Fig. 2F. Genes encoding hallmark feature of ICE elements are highlighted. (**E**) DNA gel electrophoresis analysis of products resulting from PCR targeting the indicated genes in the AID-1 containing donor strain *B. fragilis* 638R, the recipient strain *B. fragilis* 43859, and a representative clone obtained after AID-1 transfer. tRNA-Lys PCR was performed with an outward-facing forward primer on the AID-1 ICE and a reverse primer on the chromosome 3’ of the tRNA-Lys gene. (**F**) Outcomes of in vitro growth competition assays between *B. fragilis* 43859 or a derivative AID-1 ICE recipient and *B. fragilis* NCTC 9343. (**G**) Results of pairwise competition between the indicated *B. fragilis* strains in germ-free mice. Mice colonized with *B. fragilis* 9343 for one week prior to challenge with *B. fragilis* 43859. Sample collection and data presentation performed as in (**C**).

*B. fragilis* is typically found as a clonal population in the human gut microbiome, and recent studies suggest that this is in part due to active strain exclusion via the T6SS (*6, 16*). However, in gnotobiotic mouse colonization experiments, certain *B. fragilis* strain pairs inexplicably co-exist (*8*). We noted that one such pair corresponds to *B. fragilis* 9343 and *B. fragilis* 638R, the latter of which contains an AID-1 system harboring homologs of i6 and i7. To determine whether our *in vitro* results with these strains extend to a more physiological setting, we measured the fitness contribution of the orphan immunity genes encoded by *B. fragilis* 638R following pre-colonization of germ-free mice with *B. fragilis* 9343 (Fig. 3C, fig. S3D, E). Our results indicated that the cumulative protection afforded by orphan i6 and i7 genes underlies the capacity of *B. fragilis* 638R to persist during T6S-mediated antagonism *in vivo.*

Interestingly, we found that AID-1 resides on a predicted mobile integrative and conjugative element (ICE), potentially providing an explanation for its distribution (Fig. 3D) (*17*). To test whether this element can be transferred between strains, we performed mobilization studies using *B. fragilis* 638R as a donor and *B. fragilis* 43859 as a recipient. An antibiotic resistance marker was inserted within AID-1 to facilitate the detection of its transfer. With this tool, we readily detected AID-1 transfer (Fig. 3E). This occurred at a frequency of approximately 5×10^−6^, in line with previous quantification of ICE mobility in *Bacteroides* spp. (*18*). Next, we asked whether the transfer of AID-1 to *B. fragilis* 43859 is sufficient to confer resistance to T6S-mediated antagonism. *In vitro* growth competition assays against *B. fragilis* 9343 showed that AID-1 effectively neutralizes intoxication by e6 and e7 (Fig. 3F). The receipt of AID-1 also granted significant protection to *B. fragilis* 43859 against T6-mediated killing in germ-free mice pre-colonized with *B. fragilis* 9343 (Fig. 3G). Together, these findings indicate that the transfer of a mobile orphan immunity island to a naïve *Bacteroides* strain is sufficient to provide defense against T6S effectors.

Given the benefit of orphan immunity genes against *B. fragilis* effectors, we hypothesized that this mechanism of inhibiting interbacterial antagonism should extend to effectors produced by other species. We previously reported evidence that *B. fragilis* is antagonized by other *Bacteroides* spp. in the human gut microbiome (*6*). In addition to the T6SS present exclusively in *B. fragilis,* this species and other *Bacteroides* spp. can possess other T6SSs with a distinct and non-overlapping repertoire of effector and immunity genes (*6, 13*). Therefore, we searched *B. fragilis* genomes for sequences homologous to the immunity genes corresponding to these systems. In 29 of the 122 available *B. fragilis* genomes, we identified apparent orphan homologs of these immunity genes grouped within gene clusters (Fig. 4A). While analogous to the AID-1 and −2 systems, these clusters have several unique characteristics including skewed GC content, conservation of a gene encoding a predicted XerD-family tyrosine recombinase, and repetitive intergenic sequences reminiscent of those present in integrons (fig. S4A-C) (*19, 20*). These often-large gene clusters, hereafter referred to as recombinase-associated AID (*r*AID-1) systems, can exceed 16 kb and contain up to 31 genes with varying degrees of homology to T6S immunity genes and predicted immunity genes associated with other interbacterial antagonism pathways (Fig. 4A, fig. S4D, E, table S6) (*21*). Using the shared characteristics of *B. fragilis r*AID-1 systems, we searched for related gene clusters across sequenced Bacteroidales genomes. We found that over half of sequenced bacteria belonging to this order possess a *r*AID-1 system (226 of 423) (Fig. 4B, tables S6 and S7). In sum, these gene clusters contain 579 unique genes, encompassing homologs of 25 Bacteroidales T6S immunity genes.

**Fig. 4.**
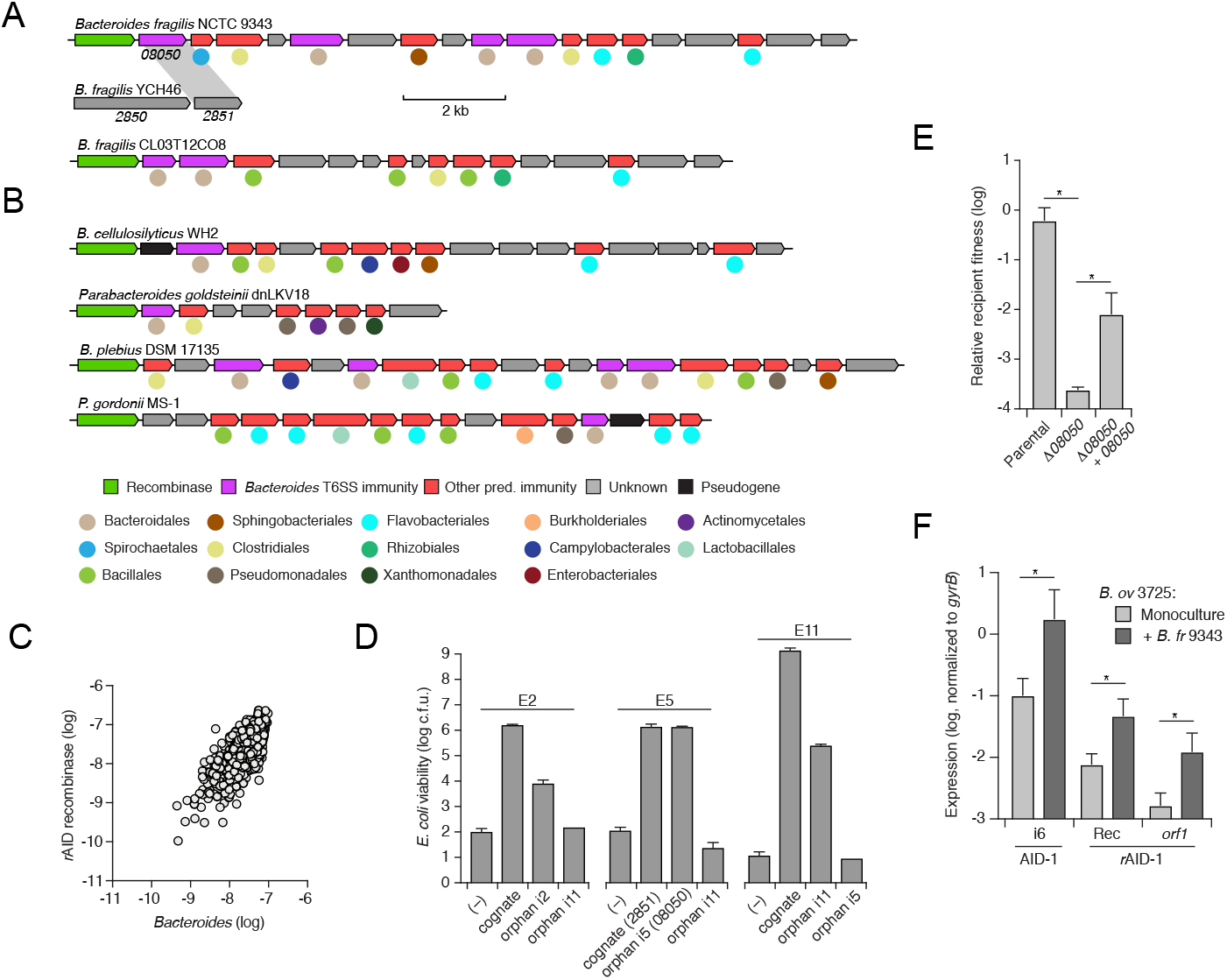
*r*AID systems encode toxin-neutralizing immunity genes and are prevalent in human gut microbiomes. (**A,B**) Depiction of *r*AID-1 clusters from the indicated *B. fragilis* (A) or Bacteroidales (**B**) species. *r*AID-1 cluster genes were assigned to functional immunity classes (indicated by gene coloring) via profile HMM scans and BLAST against a curated database of Bacteroidales T6SS immunity genes (*6, 21*). Colored circles indicate the taxonomic association of the top non-*r*AID-1 gene homolog identified via translated BLAST against the NCBI nr database. Homology (70% amino acid identity) between the first open reading frame (08050) of the *B. fragilis* 9343 *r*AID-1 cluster and a T6S cognate immunity gene from *B. fragilis* YCH46 (2851) is indicated by gray shading. (**C**) Comparison of the total abundances of *r*AID-associated predicted recombinases and the *Bacteroides* genus in adult microbiome samples derived from the HMP and MetaHIT studies (table S8). Abundance values are calculated as in Fig 1; genus abundance corresponds to the sum of all *Bacteroides* spp. (calculated individually as the average of species-specific marker gene abundances). (**D**) Viable *E. coli* cells recovered from cultures carrying plasmids expressing the indicated toxin and immunity proteins. (**E**) Outcomes of *in vitro* two-strain growth competition assays between the indicated *Bacteroides* strains. The relevant *r*AID-1 gene of *B. fragilis* 9343 and its corresponding effector within *B. fragilis* YCH46 are depicted in (**A**). (**F**) Results of qRT-PCR analyses for the indicated *B. ovatus* 3725 genes belonging to AID-1 (i6, M088_1971) or *r*AID-1 clusters (Rec, recombinase, M088_1401; *orf1*, M088_1400) under conditions of growth in mono- or co-culture with *B. fragilis* 9343 for 2 hrs.

The prevalence of *r*AID-1 genes in Bacteroidales genomes suggested that these elements may be common in the human gut microbiome. To investigate this, we searched metagenomic data for sequences mapping to Bacteroidales T6S orphan immunity genes found within *r*AID-1 systems. Remarkably, we found one or more *r*AID-1 immunity genes in 551 of 553 samples using a 97% sequence identity threshold to map reads (table S8). These *r*AID-1 immunity genes diverge significantly from corresponding cognate immunity genes (corresponding to 32-91% amino acid identity), suggesting that the latter are an unlikely source of significant false positives in this analysis. We also searched the same samples for *r*AID-1-associated recombinase sequences. Although recombinase genes are widely distributed across bacteria, close homologs (>50% amino acid identity) of those found associated with *r*AID-1 systems are restricted to this context and only found in Bacteroidales genomes. Consistent with this, we found the abundance of *r*AID-1 recombinase genes correlates strongly with the genus *Bacteroides* (Fig. 4C).

Orphan immunity genes encoded within *r*AID-1 clusters diverge more significantly from cognate immunity than do those within AID systems. Thus, we sought to experimentally validate the capacity of *r*AID-1 immunity genes to protect bacteria from intoxication. Since most bacteria harboring *r*AID-1 systems have limited genetic tools, we employed *E. coli* to identify three Bacteroidales T6S effector genes that intoxicate cells in a manner that is neutralized by cognate immunity. In each case, we found that co-expression of these effector genes with corresponding *r*AID-1-associated orphan immunity genes, but not mismatched orphan immunity genes, restored *E. coli* growth (Fig. 4D). Both genes from one effector–orphan immunity pair we validated derive from genetically tractable strains: *B. fragilis* YCH46 (effector, 2850) and *B. fragilis* 9343 (orphan immunity, 08050). *In vitro* growth competition experiments with these strains, and mutant and genetically complemented derivatives, showed that an endogenous *r*AID-1 orphan immunity gene of *B. fragilis* 9343 can neutralize a T6S-delivered toxin (Fig. 4E).

The orphan immunity systems we defined consist of many genes and their expression could incur a substantial metabolic burden. As a first step toward understanding the regulation of AID systems, we performed quantitative RT-PCR to compare the expression of the systems in the presence and absence of a competitor strain. These studies provided evidence that transcription of both systems is induced by co-cultivation with a competitor strain (Fig. 4F). We also examined metatranscriptomic data for evidence of AID expression (*22*). Owing to a paucity of such data available for samples definitively containing AID-1 and AID-2, we could not systematically quantify expression of these systems. However, using conservative criteria for defining *r*AID-1-associated genes in metatranscriptomic data, we found evidence of the expression of this system in every sample derived from a large study (n=156) (table S9). In some samples, such as those with high levels of *Bacteroides, r*AID-1 genes accounted for nearly 1/10,000 of all metatranscriptomic reads. Taken together with our functional characterization of AID systems, these findings suggest that acquisition and maintenance of consolidated orphan immunity determinants is a common mechanism by which Bacteroidales defend against interbacterial antagonism in the human gut microbiome.

Mounting evidence suggests that competitive interactions between bacteria predominate in many environments (*23*). This evolutionary pressure has undoubtedly led to the wide dissemination of idiosyncratically orphaned immunity genes predicted to provide resistance to diverse antagonistic pathways (*21, 24–27*). Modeling studies predict that interbacterial antagonism is a critical contributor to the maintenance of a stable gut community (*28*). Our findings reveal that a corollary of the pervasiveness of antagonistic mechanisms is strong selective pressure for genes that can provide protection against attack, establishing a molecular arms race that has led to diversification and expansion of T6S effectors. Deciphering the linkage between orphan immunity genes and the bacteria harboring the cognate effectors has the promise of providing a window into the physical connectivity of bacteria in the gut microbiome.

It is now appreciated that phage defense mechanisms, including the adaptive system, CRISPR-Cas, are critical for bacteria to cope with the omnipresent threat and deleterious outcome of phage infection (*3*). However, the ubiquity of interbacterial antagonistic systems suggests that in most habitats, bacteria are equally, or perhaps more likely to be subject to attack and potential cell death via the action of other bacteria (*21, 29–31*). Despite this, to date, consolidated interbacterial defense systems have not been investigated experimentally, nor are their ecological consequences understood. Our characterization of AID systems encoded by prevalent members of the human gut microbiota appears to reconcile these observations and demonstrate that the neutralization of contact-dependent interbacterial antagonism can be a critical mechanism for survival in polymicrobial environments. Additionally, it suggests that the immune systems of bacteria, in a manner analogous to those of vertebrates, includes arms specialized in viral or bacterial defense.

## Methods

### Microbiome Data

Metagenomic data from healthy adults were obtained from a number of large-scale sequencing projects. We specifically obtained 147 samples from the Human Microbiome Project 1.0, 100 samples from HMP 1.2, and 99 and 207 samples from two different MetaHIT datasets (*10–12, 32*). We further obtained paired metagenomic-metatranscriptomic data from a study of 156 individuals (*22*). Finally, we obtained a database of genes identified from 1267 assembled metagenomes as part of the Integrated Gene Catalogue (IGC) (*33*).

### Analysis of gene and species abundances in microbiome samples

We previously compiled a list of T6SS immunity and effector genes (*6*). We additionally compiled a list of species-specific marker genes for all *Bacteroides* species obtained from MetaPhlAn 2.0 (*34*). In order to determine the abundance of a given immunity, effector, or marker gene in each metagenomic sample, single end metagenomic reads were aligned to gene sequences using bowtie2, allowing for one mismatch in the seed (*35*). We counted the number of reads that aligned to each such gene with at least 80% nucleotide identity (to encompass divergent orphan immunity gene sequences) and minimum mapping quality of 20. The abundance of a gene was calculated as the number of reads aligned to this gene, normalized by the gene length and by the library size. For each species, the average gene level abundance of all species-specific marker genes was used to assess the species abundance. For the total *Bacteroides* abundance, we used the sum of all species-specific marker genes in the genus. Samples were only included in an analysis if they had at least 10 reads mapping to the type 6 genes in question (effectors, immunity or recombinases). Based on the abundance of GA3 immunity genes and *B. fragilis* we split samples into those where *B. fragilis* was not detectable, those where the immunity gene had >10X the *B. fragilis* marker gene abundance, and those where such discrepancy between the abundance of immunity genes and that of *B. fragilis* was not observed. Metatranscriptomics data was processed similarly to metagenomics data, except that abundance values were converted to an RPKM for familiarly with canonical RNA-Seq analysis.

### Orphan immunity phylogenetic analysis

Filtered reads derived from human shotgun microbiome datasets were aligned using bowtie2 as described above and subsequently converted to a pileup using samtools with parameters --excl-flags UNMAP,QCFAIL,DUP −A −q0 −C0 −B (*35, 36*). A sequence corresponding to the most abundant version of the immunity gene in the sample was reconstructed from that pileup as follows. First, 50 bases from the start and end were trimmed due to a propensity for low coverage. Second, at all sites with at least 10X coverage the base was set to the major allele. Sites with <10X coverage were assigned an ambiguous base. Finally, we only kept the reconstructed sequence in metagenomic samples where at least 90% of the sequence had >10X coverage. The number of SNPs between all immunity sequences, both from metagenomic samples and from *Bacteroides* genomes, was calculated and used to populate a distance matrix. Since obtained distances were small (e.g., a single base difference), we used hierarchical clustering (with complete linkage), rather than standard phylogenetic reconstruction methods, to visualize the relatedness between different sequences.

### Assigning orphan immunity sequences to bacterial species

We aimed to identify the species most likely to encode the immunity gene in each cluster of identical sequences reconstructed from metagenomes. Only clusters with at least 3 sequences were used, to ensure statistical confidence. The abundance of each species was assessed based on species-specific marker genes as described above. We specifically employed a simple linear model that assumed that only a single species encodes the immunity gene. We further assumed a one to one relationship between species marker gene abundance and orphan immunity gene abundance, and accordingly fixed the intercept at zero and allowed a single species with a slope of one. The fit of the model for each species was calculated as the mean square error over all samples. The most likely species to encode the immunity gene was determined by the minimum mean squared error.

### Assembly of orphan immunity sequences from metagenomes

Paired-end metagenomic sequencing data was assembled using SoapDeNovo2 with a kmer length of 63 and an average insert size of 200 (*37*). BLAST was used to identify the contig that contained the orphan immunity gene, and GeneMarkS was used to predict protein coding genes (*38*).

### Bacterial culture conditions

Anaerobic culturing procedures were performed either in an anaerobic chamber (Coy Laboratory Products) filled with 70% N_2_, 20% CO_2_ and 10% H_2_, or in Becton Dickson BBL EZ GasPak chambers. *E. coli* EC100D *λ* pir and S17-1 *λ* pir strains were grown aerobically at 37°C on lysogeny broth agar. Unless otherwise noted, *Bacteroides* strains were cultured under anaerobic conditions on brain heart infusion agar (BHI; Becton Dickinson) supplemented with 50 μg mL^−1^ hemin and 10% sodium bicarbonate (*39*). Antibiotics and chemicals were added to media as needed at the following concentrations: trimethoprim 50 μg mL^−1^, carbenicillin 150 μg mL^−1^, gentamicin 15 μg mL^−1^ (*E. coli*), gentamicin 60 μg mL^−1^ (*Bacteroides*), erythromycin 12.5 μg mL^−1^, tetracycline 6 μg mL^−1^, chloramphenicol 12 μg mL^−1^, floxuridine (FUdR) 200 μg mL^−1^.

### Genetic techniques

Standard molecular procedures were employed for creation, maintenance and *E. coli* transformation of plasmids. All primers used in this study were synthesized by Integrated DNA Technologies (IDT). Phusion polymerase, restriction enzymes, T4 DNA ligase, and Gibson Assembly Reagent were obtained from New England Biolabs (NEB). A comprehensive list of primers, plasmids, and strains are provided (Supplemental Data Table 10). Deletion of the gene encoding thymidine kinase in *B. fragilis*, *B. ovatus*, and *B. xylanisolvens* strains was performed by cloning respective genomic flanking regions into the vector pKNOCK as previously described (*40*). Briefly, pKNOCK*-tdk* plasmids were mobilized into *Bacteroides* strains via overnight aerobic mating with *E. coli*. Integrants were isolated by plating on selective media, were passaged once without antibiotics to allow for plasmid recombination, and plated for counter selection on FUdR. Recovered single colonies were patched onto selective media to ensure loss of pKNOCK, and disruption of *tdk* confirmed by PCR. Subsequent deletion of orphan immunity genes was performed in *tdk* strains via a similar counter selection strategy, except employing the suicide plasmid pExchange in place of pKNOCK (*7*). Genomic deletions were confirmed by PCR. Gene complementation was performed by cloning genes into pNBU2-erm_us1311 for constitutive expression (*41*).

### Isolation of *Bacteroides* strains from fecal samples

Fecal samples from healthy infants used for strain isolation were collected as part of a prior study approved by the Seattle Children’s Hospital Institutional Review Board (*14, 42*). Frozen stool samples stored at −80°C were manually homogenized, serially diluted in tryptone yeast glucose (TYG) broth, and plated under anaerobic conditions on *Bacteroides* bile esculin (BBE) agar plates (Oxyrase, Inc). Single colonies which exhibited esculin hydrolysis as indicated by the production of black pigment on BBE agar were sub-cultured in TYG broth with the addition of 60ug/mL gentamicin until stationary phase and then were frozen at −80°C following the addition of sterile glycerol to 20% final concentration. Single colonies isolated from these stocks were subsequently screened by PCR with primers targeting the orphan i6 gene as assembled from metagenomic short read sequence data (*14*).

### Genome sequencing

Genomic DNA used for Illumina sequencing was prepared by harvesting *Bacteroides* strains grown overnight on BHIS blood agar plates. Cells resuspended from plates were washed in phosphate buffered saline before DNA extraction with the Qiagen DNeasy Blood and Tissue Kit. Sequencing was performed on an Illumina MiSeq using the V3 Reagent kit at the Northwest Genomics Center sequencing facility at the University of Washington. Assembly was done with paired end Illumina reads using SPAdes version 3.7.1 in ‘careful’ mode (*43*). We used a 93 K-mer setting for *B. xylanisolvens* H207 and 99 K-mer setting for *B. ovatus* 3725 D1 iv. Open reading frames on genome scaffolds were annotated using Prokka 1.12 (*44*). AID clusters often appear in highly repetitive genomic contexts (e.g. mobile elements) and are often split into multiple scaffolds in reference genomes. To compensate for this, we additionally performed long read sequencing via PacBio on a subset of genomes. To this end, high-molecular weight DNA was extracted using the Qiagen Genomic-tip Kit and sequenced by SNPsaurus (Eugene, OR) using a PacBio Sequel. PacBio assemblies were generated using Canu 1.7 (*45*). Species identification was performed by blast searches with species-specific marker genes (*34*). Whole genome sequencing data generated in the course of this study has been deposited at the Sequence Read Archive under BioProject Accession PRJNA484981.

### Interbacterial competition assays

*Bacteroidales* strains were grown on BHIS blood agar plates overnight at 37°C. Bacteria were resuspended from plates in BHIS broth and the optical density of each strain was adjusted to a 10:1 *B. fragilis* NCTC 9343 to competitor ratio (OD600 6.0 to 0.6) for competitions involving *B. xylanisolvens* and *B. ovatus,* or 1:1 ratio for competitions involving *B. fragilis* 638R (OD600 6.0). Equal volumes of each strain at the adjusted OD were mixed and 5ul of bacterial mixtures were spotted onto predried BHIS blood agar plates, in triplicate spots. Competitions were allowed to proceed for 20-24 hours at 37°C under anaerobic conditions before spots were harvested into BHIS broth. Competition outcomes were quantified in one of two ways: 1) by serial dilution for enumeration of colony forming units after plating on BHIS selective plates containing either erythromycin or tetracycline, or 2) purification of total genomic DNA using the Qiagen DNeasy Blood and Tissue Kit and subsequent quantification by qPCR using strain-specific primers (see Supplemental Data Table 6). For antibiotic selection, *B. fragilis* 9343 was marked with erythromycin resistance by integration of pNBU2-erm at the att1 site (*41*). Other strains were either naturally tetracycline resistant, or were marked by integration of pNBU2-tet-BCO1. Strains with insertions of pNBU2 were selected for matching integration sites by PCR with primers flanking att loci (*46*). Interbacterial competitions between strains of *B. fragilis* occasionally exhibited T6SS-independent phenotypes that were dependent on the initial starting ratio of the strains used (*47*).

### Interbacterial mobile element transfer assays

Allelic exchange was used to engineer a high-expression chloramphenicol resistance cassette onto the AID-1 system of *B. fragilis* 638R, replacing BF638R_2056-2058 (*48*). Chloramphenicol-resistant *B. fragilis* 638R cells were mixed on BHIS blood agar plates with erythromycin-resistant *B. fragilis* ATCC 43859 cells at a 1:1 ratio (OD600 6.0). Following overnight co-culture, bacterial mixtures were harvested and plated on BHIS plates containing either erythromycin alone (to quantify c.f.u. of total ATCC cells), or erythromycin and chloramphenicol (to quantify c.f.u of AID-1 recipient ATCC cells). Doubly-resistant colonies were screened individually by PCR to confirm strain identity, the presence of the AID-1 system, and the genomic integration site at a tRNA-Lys gene (see Table S10 for primers used).

### Gnotobiotic animal studies

Germ-free 6-12 week-old female Swiss Webster mice from multiple litters were randomized, housed simultaneously in pairs in single Techniplast cages with a 12-hour light/dark cycle, and fed a standard lab diet (Laboratory Autoclavable Rodent Diet 5010, LabDiet), in accordance with guidelines approved by the University of Washington Institutional Animal Care and Use Committee. Blinding was not performed. *Bacteroides fragilis* strains were introduced into mice via oral gavage of 10^8^ colony forming units (c.f.u.) suspended in 0.2mL of sterile 1X phosphate buffered saline with 20% glycerol. Challenge with *B. fragilis* 638R or *B. fragilis* ATCC strains occurred 7 days following pre-colonization with *B. fragilis* 9343 strains. Colonization levels by each strain in each mouse were tracked by collection of fecal pellets over a period of 4 weeks, plating on selective BHIS agar plates (*B. fragilis* 9343 on BHIS plus erythromycin; *B. fragilis* 638R and ATCC on BHIS plus tetracycline), and subsequent absolute quantification of c.f.u. by normalization of each sample to the initial pellet weight. Differences in the strain ratio of c.f.u between groups at each timepoint was assessed using Mann Whitney U tests. Non-parametric tests were used following Shapiro-Wilk analysis for normality of data at each time point. Mice were confirmed to be sterile prior to colonization by qPCR with primers targeting the 16S rRNA gene and free of non-*Bacteroides* contamination by plating fecal pellets on non-selective LB and BHIS plates incubated under either anaerobic and aerobic conditions (*49*).

### Bioinformatic analysis of *r*AID clusters

The amino acid sequence of the *B. fragilis* NCTC 9343 polyimmunity-associated XerD-like tyrosine recombinase (BF9343_RS08045) was used as a query against a custom database of 423 *Bacteroidales* genomes downloaded from GenBank. rAID clusters in *Bacteroidales* genomes were identified based upon the following criteria: i) presence of a 5’ XerD-like tyrosine recombinase gene encoding a protein with amino acid identity exceeding 44% (corresponding to an e-value of 10^−100^), ii) 2 or more co-directionally oriented downstream genes which possessed iii) a GC content of 41% or lower. The end of the gene cluster was defined as the stop codon of the last co-directionally oriented gene in the cluster with similar GC content. To identify homologs of genes within rAID clusters, open reading frames within the clusters were translated and used as tBlastn queries against the NCBI non-redundant nucleotide database. Top hits from these searches were often genes in other rAID clusters; therefore, these hits were discarded. The top non-*r*AID hit from tBlastn searches with an e-value threshold of 10^−30^ was selected as the closest homolog. *r*AID cluster genes were assigned to interbacterial immunity gene families via hmm scans with profiles from Zhang et al (*21*) with an e-value cutoff of 10^−3^. rAID cluster genes were additionally compared via tblastn with forty-six Bacteroidales T6SS immunity genes from subtypes 1-3 (*6, 13*) with an e-value cutoff of 10^−10^. Percent amino acid identity with homologs was assessed if sequences could be aligned across ≥80% of their length. Motif enrichment analysis was performed on noncoding sequences within a subset of *r*AID-1 clusters (14 sequences immediately 3’ of the recombinase stop codon, and 86 intergenic sequences between *r*AID-1 ORFs), using MEME Suite 5.0.2 and default settings (*50*).

### Heterologous expression of *Bacteroides* toxin and immunity genes

To assess the ability of cognate immunity or orphan immunity to neutralize the toxicity of a *Bacteroidales* T6SS effector, genes were cloned into *E. coli* expression vectors pScrhab2-V (effectors) and pPSV39-CV (immunity). Immunity genes were fused with the *P. aeruginosa* ribosome binding site from *hcp1* during the cloning process (*51*). All cloning steps for effector genes involved growth of *E. coli* on media containing 0.1% glucose to ensure repression of expression. *E. coli* DH5a cells were co-transformed with pSchraB2 and pPSV39 plasmids bearing genes of interest. Overnight cultures were then grown from single co-transformed colonies to stationary phase in LB broth containing 50ug/mL trimethoprim, 15ug/mL gentamycin, and glucose. Cells were harvested from these cultures and washed to remove glucose before back-dilution to an OD600 of 0.05 into LB broth containing 50ug/mL trimethoprim, 15ug/mL gentamycin, and either no inducer, 0.05% rhamnose only, or 0.05% rhamnose and 1mM isopropyl β-D-1-thiogalatopyranoside (*51, 52*). Cultures were then grown for 8 hours shaking at 37³C before plating to allow quantification of c.f.u.

### Gene expression analysis of AID-1 and *r*AID-1 systems of *B. ovatus* 3725

To assess the level of expression of genes in the AID-1 and *r*AID-1 systems of *B. ovatus* 3725, bacterial cells were first grown overnight on BHIS blood agar plates containing gentamycin. Cells were then resuspended in BHIS to an OD600 of 3.0 for *B. ovatus* monocultures, or to an OD600 of 0.3 for mixed co-culture experiments with *B. fragilis* 9343 at 10-fold excess (OD600 of 3.0). 5uL volumes of bacterial mixtures were then spotted on BHIS blood agar plates. Plates were incubated at 37³C for 2 hours under anaerobic culture conditions before cells were harvested directly in Buffer RLT plus (twenty 5uL spots per condition per replicate, Qiagen RNeasy Micro Kit). Two separate rounds of DNase treatment were performed (Qiagen RNase-free DNase, Thermo Fisher Scientific Turbo DNase-free kit). RNA samples were confirmed to be free of genomic DNA by PCR with primers targeting the Bacteroides 16S rRNA gene. cDNA was generated using the High Capacity cDNA Reverse Transcription Kit (Applied Biosciences). Following synthesis, cDNA was diluted 1:10. Quantitative PCR (primers listed in Table S10) was performed using SSO Universal SYBR Green Supermix (Bio-Rad) on a CFX96 machine (Bio-Rad). Genomic DNA was used to generated standard curves (*53*). Differences in gene expression between samples was performed by normalization to the expression level of *B. ovatus* 3725 *gyrB.* Primers targeting *gyrB* were designed to target regions of the genes that are highly polymorphic between *B. fragilis* and *B. ovatus*, and species-specificity for *B. ovatus* was confirmed by PCR using *B. fragilis* genomic DNA (*54*).

## Supporting information

19_Ross_bioRxiv_SupplementalTables1-10

## Acknowledgements

We thank the UW GNAC for assistance with gnotobiotic experiments. We thank Cynthia Sears, Andy Goodman, Tomomi Kuwahara, and Eric Martens for generously providing *Bacteroides* strains. Funding: This work was supported by National Institutes of Health grants AI080609 (to JDM), P30DK089507 (to LRH as pilot study PI), R01DK095869 (to LRH), K99GM129874 (to BDR), R01GM124312 (to EB), and New Innovator Award DP2AT00780201 (to EB), and the Burroughs Wellcome Fund (to JDM). AJV was supported by a postdoctoral fellowship from the Natural Sciences and Engineering Research Council of Canada. BDR was supported by a Simons Foundation-sponsored Life Sciences Research Foundation postdoctoral fellowship. EB is a Faculty Fellow of the Edmond J. Safra Center for Bioinformatics at Tel Aviv University. JDM is an HHMI Investigator.

## Author Contributions

BDR, AJV, AHM, EB, and JDM designed the study. BDR, AJV, MCR, DTS, CEP, and LRH performed experiments. BDR, AJV, MCR, DTS, AHM, SBP, EB, and JDM analyzed data. BDR, AJV, SBP, EB, and JDM wrote the manuscript.

## Data and materials availability

*Bacteroides* strains acquired from Johns Hopkins University were obtained under an MTA. All data required to assess the conclusion of this research is available in the main text and supplemental materials, or has been deposited at the Sequence Read Archive under BioProject Accession PRJNA484981. Analysis code is available upon request.

## Supplemental Tables

**Table S1.** Metagenomic results derived from the HMP and MetaHIT studies utilized in this study.

**Table S2.** List of Bacteroidales T6SS cognate immunity genes.

**Table S3.** Accession numbers of *Bacteroides* genomes harboring orphan i6 genes.

**Table S4.** Metagenomic results derived from infant stool samples utilized in this study.

**Table S5.** Description of AID gene clusters in genomes depicted in Fig. 2.

**Table S6.** Description of *r*AID-1 gene clusters depicted in Fig. 4.

**Table S7.** Features associated with *r*AID-1 gene clusters in Bacteroidales genomes.

**Table S8.** Metagenomic results of *r*AID-1 genes derived from the HMP and MetaHit studies utilized in this study.

**Table S9.** Metagenomic and metatranscriptomic results from analysis of *r*AID-1 genes from PMID 29335555.

**Table S10.** Strains, plasmids and primers used in this study.

---

**Fig. S1.**
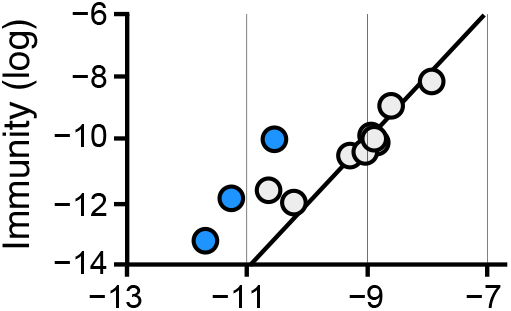
Prevalence of *B. fragilis*-specific orphan immunity genes in infant microbiomes. Comparison of the abundance of *B. fragilis-specific* T6SS immunity genes with *B. fragilis* species-specific marker genes in infant microbiome samples (Table S4) (*14*). Abundances are calculated as in Fig. 1A. Samples in which immunity gene abundance exceeds that of *Bacteroides* by over 10-fold (blue) are highlighted.

**Fig. S2.**
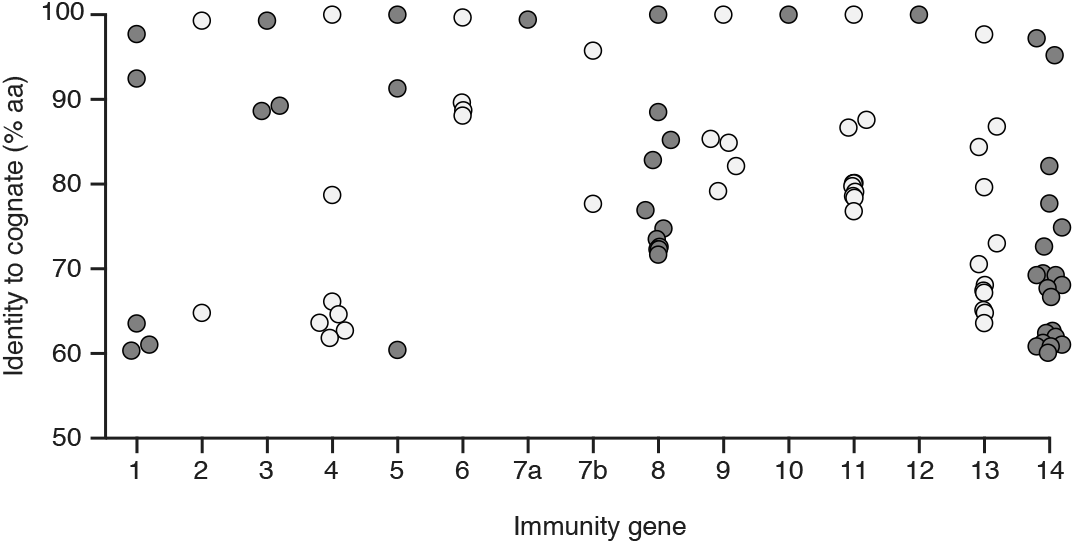
Diversity and genomic context of orphan immunity genes in human gut microbiomes. Data points indicate the amino acid identity of unique genes homologous to indicated *B. fragilis-specific* T6SS cognate immunity genes identified through BLAST analysis of the Integrated Gene Catalog (IGC) (maximum E-value, 10^−40^; minimum percent identity, 60%) (*33*).

**Fig. S3.**
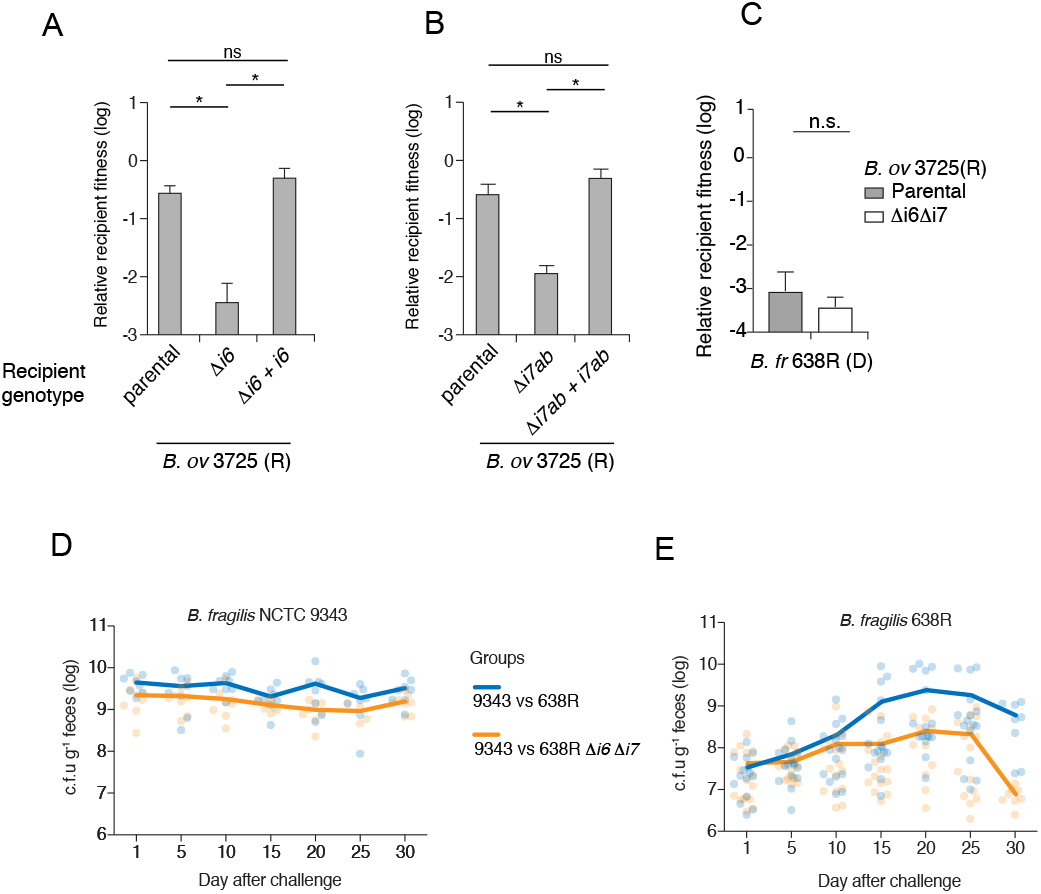
Orphan immunity genes specifically enhance the fitness of *Bacteroides* strains *in vitro* and *in vivo.* (**A,B**) T6SS-dependent competitiveness of parental strains of *B. ovatus* 3725 and the indicated mutant and complemented derivatives during *in vitro* growth competition experiments with *B. fragilis* 9343. Relative recipient fitness was determined by calculating the ratio of final to initial colony forming units and normalizing to the corresponding experiment with *B. fragilis* 9343 lacking *tssC* (T6S-inactive). Data represent mean ± s.d. of three replicates. Asterisks indicate statistically significant differences between indicated mean values (t-test, p < 0.01). (**C**) T6SS-dependent competitiveness of a parental strain of *B. ovatus* 3725 or a strain bearing inframe deletions of indicated orphan immunity genes, during *in vitro* growth competition experiments with a *B. fragilis* 638R parental strain or a strain lacking *tssC* (T6-inactive). Relative recipient fitness and statistics were calculated as in (**A,B**). (**D,E**) Recovery of *B. fragilis* 9343 (**D**) or 638R and the indicated orphan immunity mutant derivative (**E**) from pairwise competitions of the strains in germ-free mice. Alternating time points of these data are included in ratio form in Fig. 3C.

**Fig. S4.**
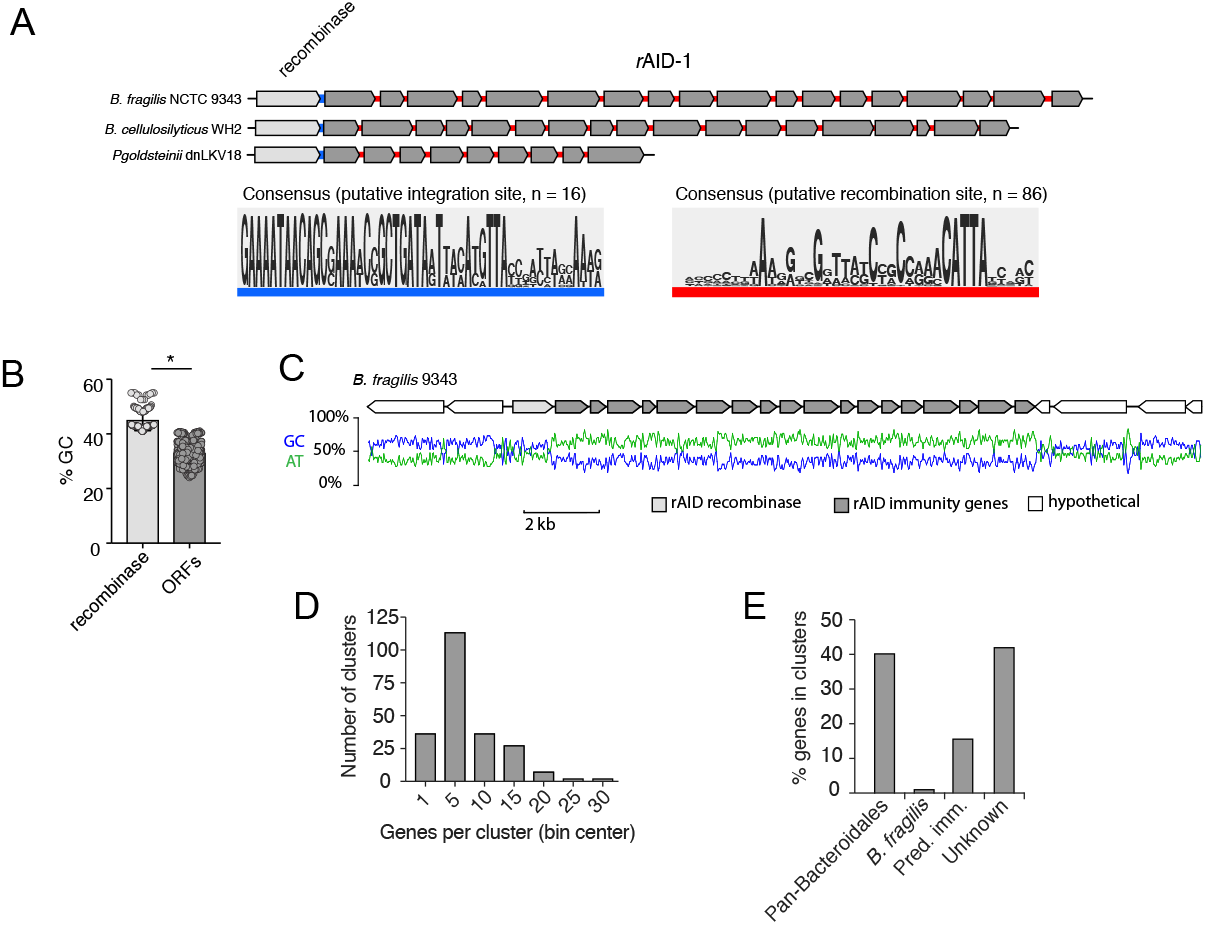
*r*AID-1 systems share hallmarks of integrons and horizontal gene transfer. (**A**) Left - Motif enrichment analysis from the intergenic sequences immediately 3’ of the recombinase stop codon to the start codon of the first downstream open reading frame within 16 randomly selected rAID-1 gene clusters. This region is highlighted in blue in three representative rAID-1 systems shown above. Right - Motif enrichment analysis from all 86 intergenic sequences between the ORFs of six *r*AID-1 clusters (*B. fragilis* NCTC 9343, *B. cellulosilyticus* WH2, *B. ovatus* 3725, *Paraprevotella clara* YIT 11840, *Parabacteroides goldsteinii* dnLKV18, and *Parabacteroides gordonii* MS-1) (*50*). This region is highlighted in red in three representative rAID-1 systems shown above. (**B**) Average G+C nucleotide content of rAID-1-associated recombinase versus *r*AID-1 predicted ORFs (*n* = 226). (**C**) Schematic depicting the G+C and A+T nucleotide content across a representative *r*AID-1 system from *B. fragilis* 9343. (**D**) Frequency distribution of gene number in *r*AID-1 clusters (*n* = 1247 genes in 226 clusters). Bin width is 5 genes. (**E**) Composition of genes in *r*AID-1 clusters (*n* = 226 clusters) as determined by profile HMM scans and BLAST analysis against a curated database of Bacteroidales T6SS immunity genes (*6, 21*).

